# Centromeres in budding yeasts are conserved in chromosomal location but not in structure

**DOI:** 10.1101/2025.07.24.666568

**Authors:** Conor Hession, Kevin P. Byrne, Kenneth H. Wolfe, Geraldine Butler

## Abstract

The budding yeast *Saccharomyces cerevisiae* has ‘point’ centromeres, which are much smaller and simpler than centromeres of most other eukaryotes and have a defined DNA sequence. Other yeast taxa have different and highly diverse centromere structures, but a clear picture of how yeast centromeres have evolved is lacking. Here, we investigated nine yeast species in two taxonomic orders that are close outgroups to *S. cerevisiae*. We find that they have a wide diversity of centromere structures, indicating that multiple transitions of structure have occurred within the last 200 Myr. Some species have centromeres with defined sequence motifs (17 – 200 bp), others consist of Inverted Repeats (IRs), and others have Ty5-like retroelement clusters. Strikingly, the chromosomal locations of centromeres have largely been conserved across taxonomic orders, even as their structures have changed, which suggests that structure replacement occurs *in situ*. In some *Barnettozyma* species we find that a single genome can contain chromosomes with different centromere structures – some with IRs and some without – which suggests that a structural transition is underway in this genus. We identified only one example of a centromere moving by a long distance: a new centromere formed recently at the *MAT* locus of *Barnettozyma californica*, 250 kb from the previous centromere on that chromosome.

**Author summary:** Centromeres are an evolutionary paradox. Their molecular function is highly conserved, but their structures vary tremendously among eukaryotes. The “point” centromeres of the yeast *Saccharomyces cerevisiae* are among the most unusual: they are tiny (< 200 bp) and non-repetitive, and unlike other centromeres they contain a single copy of a well-defined sequence motif. However, we have little knowledge about where these point centromeres came from, or more generally about how changes of centromere structure occur during evolution. Here, we characterized centromere structure and location in nine species of budding yeasts, spanning an evolutionary depth of approximately 200 million years. We find that the chromosomal locations of centromeres are extraordinarily well conserved, whereas their structures vary greatly. We show that sequence-defined centromeres are older and more widely distributed than previously realised. We identify some species in which the centromeres of different chromosomes have different structures, which suggests that they are in transition from one structure to another. The centromere variation observed makes it difficult to infer the ancestral structure.

## Introduction

Centromeres are the points of assembly of the kinetochore, the protein complex that connects the spindle microtubules to the chromosomes, enabling efficient and accurate separation of chromosome pairs during division of the nucleus in mitosis and meiosis. Centromeres are an evolutionary paradox: they are strongly conserved in function, but they change rapidly in sequence and structure over evolutionary time. Centromeres range from structures that cover the entire chromosome (holocentromeres), to regional centromeres of several thousand basepairs, to the short “point” centromeres of < 200 bp found in some yeast species such as *Saccharomyces cerevisiae* [1]. In many plants and animals, centromeres consist of many highly repetitive tandem arrays (satellites), which can be up to several Mbp long [2]. Other centromeres are highly enriched for retrotransposons [1]. The vast diversity of centromeres has undermined attempts to characterize their mode of evolution, and even their location and structure in some species.

Centromeres in most eukaryotes are defined by the localization of a nucleosome that contains a specific variant of histone H3, called CENP-A, CenH3, or (in budding yeasts) Cse4. Binding of CENP-A recruits other kinetochore proteins. However, CENP-A has been lost in some insect species with holocentromeres [3] and, more unusually, in many species of Mucoromycotina, an early diverging fungal lineage [4]. In many eukaryotes, centromeres lie in regions devoid of genes (though they may be transcribed), which are enriched in heterochromatin established either by methylation of histone H3 lysine 9, or by RNAi [1, 5]. However, most budding yeast species have lost the H3K9 methyltransferase [6] and some have lost the RNAi machinery [7]. Centromeres of some fungi are rich in A+T bases, and in fission yeast *Schizosaccharomyces pombe* almost any A+T-rich sequence can function as a centromere [8]. However, this again is not a universal feature, and many budding yeast species have centromeres that are not A+T-rich [9].

The budding yeasts (subphylum Saccharomycotina) represent an excellent resource to study structure, location and evolution of centromeres. There are 12 taxonomic orders containing species with diverse lifestyles [10, 11], covering an evolutionary time span of approximately 400 million years, and their genomes have been sequenced comprehensively [12, 13]. Budding yeasts are already known have an astonishing diversity of centromere structures, even though relatively few species’ centromeres have been studied in depth [5]. The first centromeres described at a molecular level from any eukaryote were those of *Saccharomyces cerevisiae*, a member of the order Saccharomycetales [14, 15]. These “point” centromeres are short, sequence-defined structures, approximately 125 bp in length, that include three Centromere Determining Elements (CDEI, CDEII and CDEIII) [16]. The CBF3 protein complex and the Cse4-chaperone protein Scm3 bind to the 25 bp CDEIII element, recruiting Cse4 to CDEII [17, 18]. Binding of proteins to CDEII, and of Cbf1 to CDEIII, introduces DNA bends. There are similar point centromeres in other Saccharomycetales species [19–21], although the sequences of the CDE elements are dramatically different in one genus, *Naumovozyma* [22]. It has been speculated that yeast point centromeres may have been derived from a natural plasmid called the 2 micron plasmid [23].

Beyond the order Saccharomycetales, centromeres have been characterized from multiple species in two other budding yeast orders – Serinales and Pichiales. They show extensive variation in structure and are substantially different from the sequence-defined centromeres of *Saccharomyces cerevisiae*. In Serinales, centromeres of *Candida albicans* and *Candida dubliniensis* are gene-free regions of 4-18 kb, with 3-5 kb occupied by Cse4, and no sequence conservation among them [24–26]. They are essentially featureless, and often described as short, regional centromeres. In contrast, some other Serinales species have centromeres whose most striking feature is either an inverted repeat (IR) formed by pair of sequences a few kilobases long (e.g. *Candida tropicalis* [27, 28]), or regions of high A+T content (e.g. *Clavispora lusitaniae* [29]), or clusters of Ty5-like retrotransposons and their LTRs (Long Terminal Repeats, e.g. *Scheffersomyces stipitis* [30, 31]). For Serinales species with IRs at their centromeres, the middle region located between the IRs on different chromosomes can be either homogenized (*Candida tropicalis* [27, 28]) or unique (*Candida parapsilosis* [32]). In the order Pichiales, the known structures include short sequence-defined centromeres (*Kuraishia capsulata*, with a 200 bp motif repeated several times on some chromosomes [33]), centromeres with IRs and no homogenization (*Komagataella phaffii* [34]), centromeres with IRs and homogenization (*Pichia kudriavzevii* [35]), and centromeres containing clusters of Ty5-like elements and LTRs but without IRs (*Ogataea polymorpha* [36, 37]). The total length of the centromere region in budding yeasts ranges from less than 200 bp in *Saccharomyces cerevisiae* to more than 30 kb in *Pichia kudriavzevii* and *Scheffersomyces stipitis* [31, 35].

The diversity of centromere structure and size in budding yeasts suggests that the sequence-defined point centromeres of *Saccharomyces cerevisiae* and other Saccharomycetales species are highly derived, raising many questions about their origin, about the ancestral state of centromeres in budding yeasts, and about the evolutionary mechanism by which centromere structures change. Here, our aim was to address these questions by characterizing the location and structure of centromeres in multiple species in the two taxonomic orders that are the next-closest relatives of Saccharomycetales: the order Saccharomycodales (containing the genus *Hanseniaspora*), which diverged from Saccharomycetales approximately 138 million years ago, and the order Phaffomycetales (containing the genera *Starmera, Wickerhamomyces, Cyberlindnera* and *Barnettozyma*), which diverged from common ancestor of Saccharomycetales and Saccharomycodales approximately 208 million years ago [13]. We find extensive changes of centromere organization, even within this relatively short time span, and we identify one genus (*Barnettozyma*) in which a transition of centromere structure appears to be underway. We show that centromeres usually change structure without changing location, although we also identify a single case in which a new centromere was formed hundreds of kilobases from the previous site. We find so many changes of centromere structure that it is still difficult to infer what type of centromere was present in the last common ancestor of subphylum Saccharomycotina.

## Results

### Hi-C contact maps for Saccharomycodales and Phaffomycetales species

We used high-throughput chromatin conformation capture (Hi-C) to map centromere locations in two Saccharomycodales species and seven Phaffomycetales species (Figure 1A). The species of Phaffomycetales were chosen to span the phylogenetic breadth of this order. It should be noted that the genus *Wickerhamomyces* is polyphyletic, with *W. canadensis* being more closely related to the genus *Barnettozyma* than it is to *W. anomalus* (Figure 1A) [12, 13, 38].

**Figure 1.**
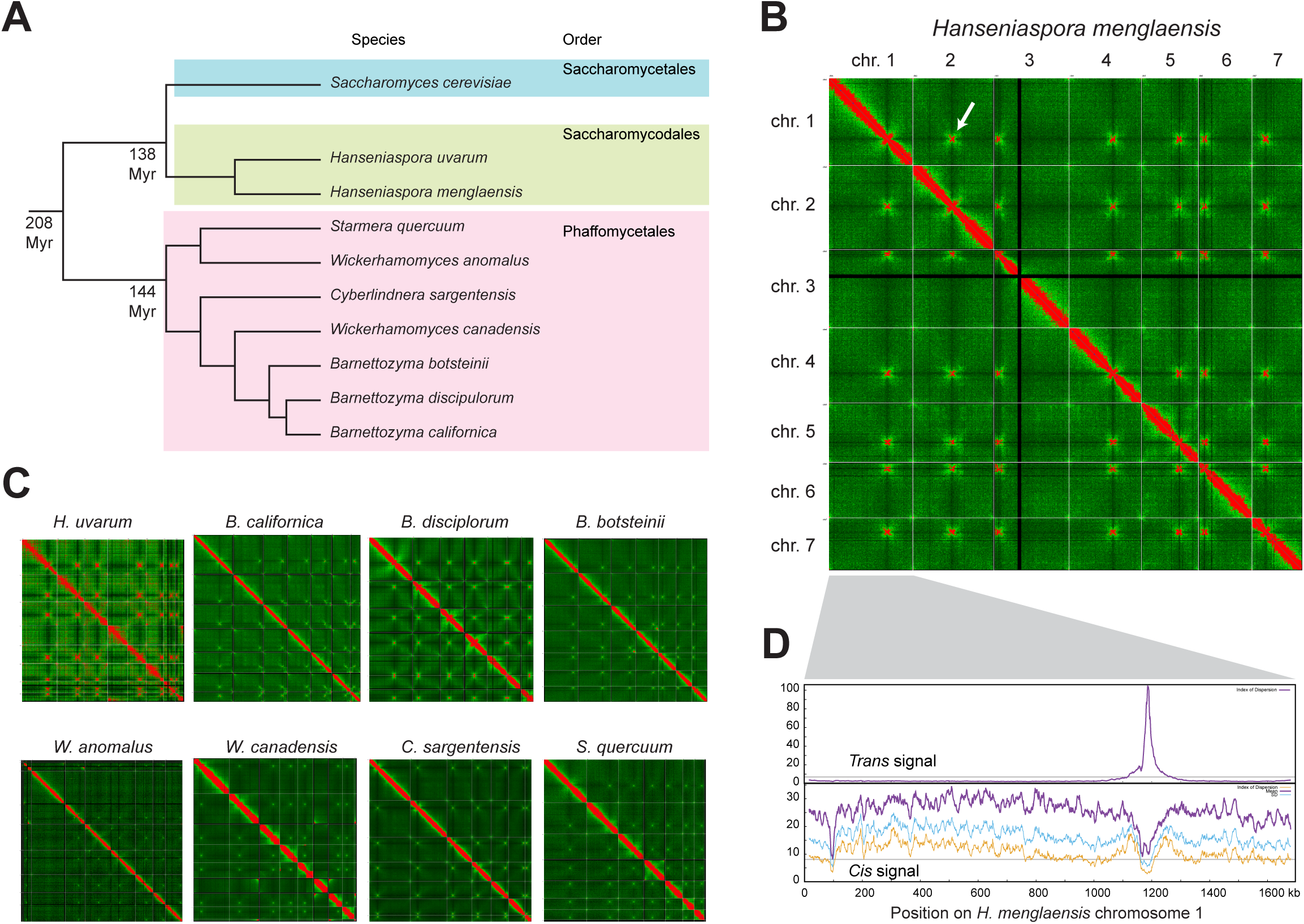
Identification of centromeres by using Hi-C. **A**. Phylogenetic tree of the yeast species studied here. Estimated divergence times between the taxonomic orders are from ref. [13]. The tree topology (not drawn to scale) is from refs [12, 38, 43]. **B**. Heatmap of Hi-C interactions in the *H. menglaensis* genome. Each pixel represents a 10 kb window. Colors were normalized by dividing the total number of interactions by the length of the genome (HiC/bp). Pixels with a HiC/bp ratio >20 are colored red, and a green gradient is used for values <20. Gray lines delineate the ends of each chromosome. The arrow highlights the Hi-C signal from the interaction between the centromeres of chromosomes 1 and 2. Similar signals are seen at the other pairs of centromeres. The black lines on chromosome 3 correspond to the rDNA locus. **C**. Hi-C heatmaps of the other species studied. **D**. Hi-C *trans* and *cis* signals on *H. menglaensis* chromosome 1. The upper panel plots the Index of Dispersion (IoD) of *trans* interactions between 10-kb windows on this chromosome and other chromosomes. The lower panel plots *cis* interactions between each 10-kb window of this chromosome and other windows on the same chromosome, showing the average number of interactions (purple), their standard deviation (blue), and their IoD (yellow).

Hi-C sequencing reads were newly generated for all species except *Hanseniaspora uvarum*, for which we utilized the raw data from a preexisting Hi-C experiment [39]. All Hi-C reads were mapped onto genome assemblies made from long-read data (Oxford Nanopore or Pacific Biosciences), either from the same isolate as used for Hi-C sequencing, or from a closely related isolate of the same species (Table S1). Four genome assemblies were newly generated for this study (Table S2), and five others were previously available [38, 40–43]. Most of the genome assemblies consist of seven complete nuclear chromosomes. *Wickerhamomyces anomalus* appears to be a hybrid species so we used the genome assembly of the primary haplotype of *W. anomalus* isolate KG16, which has 9 nuclear contigs (accession number JAHTLX01; [41]). The assemblies we used from *Hanseniaspora uvarum* and *Wickerhamomyces canadensis* also had more than 7 nuclear contigs (8 and 9, respectively; Table S1).

Hi-C contact heatmaps for the nine species are shown in Figure 1B,C. In each plot, red colors indicate the regions of the genome that are in strongest contact with each other. As shown previously for other species, as well as the major diagonal of contacts within each chromosome, there is also an inter-chromosomal signal of contacts between the centromere region of each chromosome and the centromeres of every other chromosome (e.g. the arrow in Figure 1B), which is caused by the physical proximity of all the centromeres when they are attached to the mitotic spindle during cell division [44, 45]. The heatmaps show that there are 7 centromeres, i.e. 7 chromosomes, in all nine species – even in those whose assemblies have a higher number of contigs.

### Using Hi-C cis and trans signals to locate centromeres

Each pixel in the Hi-C heatmaps (Figure 1B,C) represents the number of contacts between two 10-kb sections of the genome. To more accurately define the location of centromeres, we used a sliding window approach (window size 10 kb, step size 1 kb) and analyzed the inter-chromosomal (*trans*) and intra-chromosomal (*cis*) interactions of each chromosome separately. For each 10-kb window we calculated the number of *trans* interactions between that window and all other chromosomes, and its standard deviation, and then plotted its Index of Dispersion (IoD = StandardDeviation^2^ / Mean). Peaks in the *trans* IoD signal indicate regions on a chromosome that have high numbers of interactions with other chromosomes, such as the strong *trans* peak near position 1200 kb on *Hanseniaspora menglaensis* chromosome 1 (Figure 1D). This signal captures information from the six strong interactions that this region has with the centromere regions of the six other chromosomes (Figure 1B).

Similarly, we analyzed the *cis* interactions among windows within each chromosome, after first removing the interactions on the major diagonal. Centromeric regions tend to be depleted of *cis* interactions and appear as dips in plots of the number of *cis* interactions and their IoD, such as the trough of *cis* interactions in the same region of *H. menglaensis* chromosome 1 (Figure 1D). This signal captures information from the dark bands that are visible running horizontally and vertically through each centromeric region in the heatmaps. The *cis* and *trans* signals are independent sources of information about the centromere location on each chromosome. In general, we used the IoD peaks of *trans* interactions to define centromere locations, and referred to the *cis* interaction data for confirmation if there was any ambiguity. This approach enabled us to define the 10-kb window most likely to contain the centromere on each chromosome in each species (Table S3). Because the window step size was 1 kb, each centromere is mapped to a 10-kb interval whose endpoints have 1-kb resolution.

### Validation of Hanseniaspora uvarum centromeres using ChIP-seq

For one species, *H. uvarum*, we validated its centromere locations by using ChIP-seq to identify the binding sites of Cse4, the variant histone H3 found at yeast centromeres. A synthetic version of the *H. uvarum CSE4* gene, containing a triple HA epitope tag was cloned in plasmid pJJH3200 [46] and expressed in *H. uvarum* (Figure S1). The tag was introduced in an area of low conservation of Cse4 protein sequence among *Hanseniaspora* species. Expression of the tagged Cse4 protein was confirmed by Western blot (Figure S1). Chromatin was immunoprecipitated using anti-HA beads, sequenced, and mapped to the same genome assembly as used for the Hi-C analysis. The results showed a single strong ChIP-seq signal on each *H. uvarum* chromosome. All the ChIP-seq peaks lie in intergenic regions in *H. uvarum* (Figure 2). An independent ChIP-seq analysis of centromere locations in *H. uvarum* was very recently reported by Haase et al. [47], and our locations are in complete agreement with theirs.

**Figure 2.**
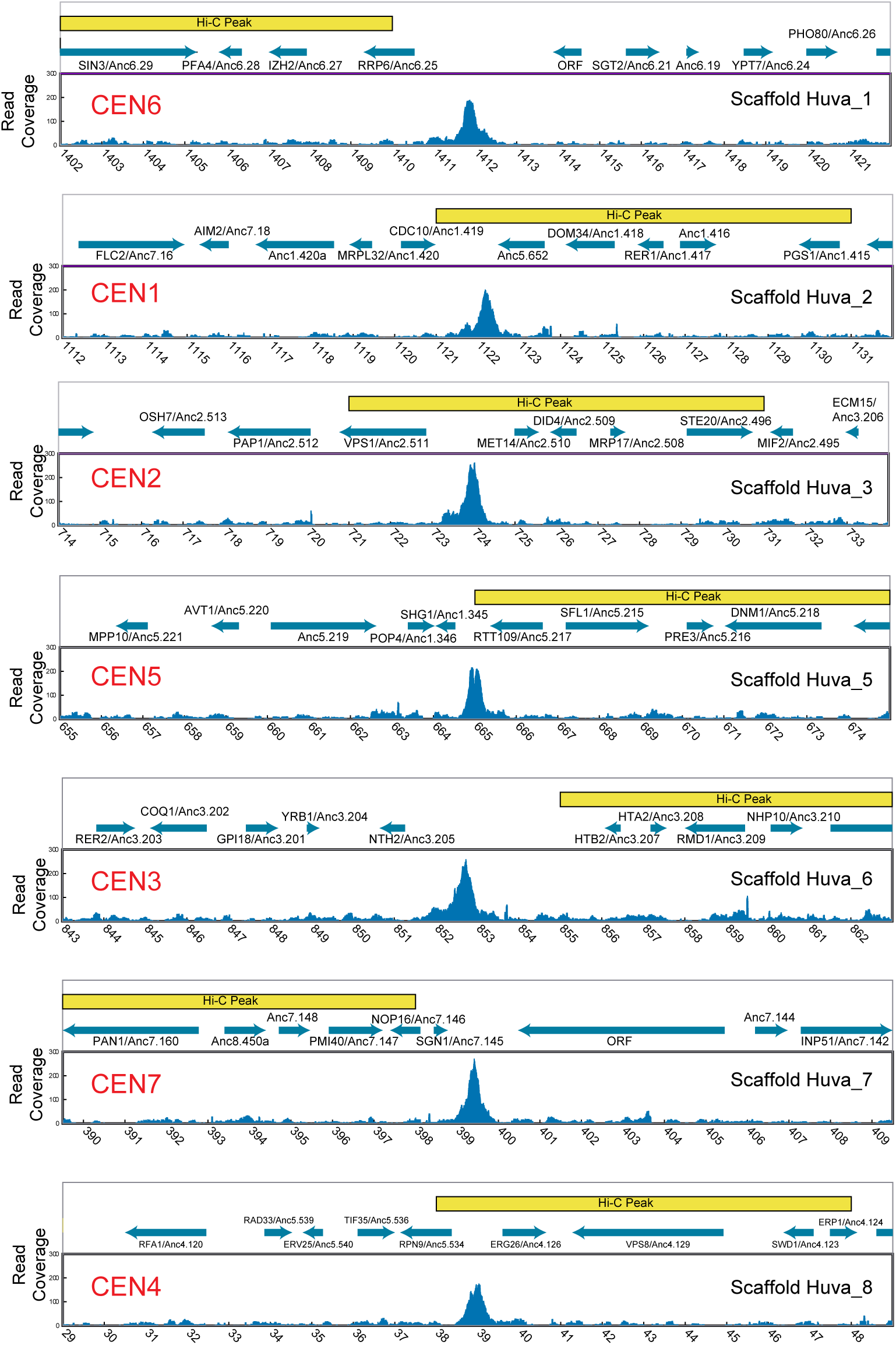
Mapping of centromeres in *H. uvarum* by ChIP-seq and Hi-C. Each panel shows the centromeric region of one *H. uvarum* chromosome, labeled with their ancestral centromere numbers (CEN, in red) and their scaffold numbers in the *H. uvarum* genome assembly (strain CBA6001; [40]). The lower part of each panel shows Cse4 occupancy (ChIP-seq read coverage) in a 20-kb window centered on the ChIP-seq peak. Yellow boxes show the 10-kb window identified as the Hi-C peak on each chromosome. Blue arrows show the locations of genes, named according to their *S. cerevisiae* orthologs and their “Anc” gene numbers in the ancestral pre-WGD Saccharomycetales genome [50].

The ChIP-seq results from *H. uvarum* allow us to gauge the accuracy of Hi-C as a method for locating centromeres (Figure 2). For four *H. uvarum* centromeres (Scaffolds 2, 3, 5, and 8), the highest ChIP-seq peak lies within the corresponding 10-kb Hi-C peak window. For the other three scaffolds, the highest ChIP-seq peak is less than 3 kb from one end of the Hi-C peak window. The Hi-C peaks that we identified in all nine yeast species therefore either contain the centromeres, or are very close to them.

### Centromere location is conserved between orders Saccharomycodales and Saccharomycetales

*S. cerevisiae* and closely related species have 16 chromosomes, resulting from a Whole Genome Duplication (WGD) [48, 49]. Most other Saccharomycetales species that did not undergo WGD have 8 chromosomes. Gordon et al. [50] reconstructed the “ancestral” gene order that existed immediately prior to the WGD event, including the location of the centromeres of the 8 ancestral chromosomes. The inferred ancestral genes are numbered with respect to chromosome and chromosomal position (e.g. Anc_1.419 is the 419^th^ gene on ancestral chromosome 1), and the centromeres are identified by ancestral chromosome number (e.g. Anc_CEN1). Centromere locations are almost completely conserved among all Saccharomycetales species and correspond to the Anc_CEN locations, except in a few taxa where the number of chromosomes has been reduced by chromosome fusion and an ancestral centromere has been lost [21, 51].

The Hi-C data from *H. menglaensis* shows that its centromere locations are generally well conserved with those in *H. uvarum* (Figure 3). In addition, the regions surrounding the 7 *Hanseniaspora* (Saccharomycodales) centromeres are syntenic with 7 of the 8 ancestral centromere locations in Saccharomycetales; they correspond to the locations of ancestral centromeres Anc_CEN1 to Anc_CEN7 so we name them CEN1 to CEN7 accordingly (Figure 3). There are some rearrangements, for example around CEN4, but overall, the conservation of gene order around centromeres between Saccharomycodales and Saccharomycetales is striking. Minor changes have occurred at CEN5 and CEN6. In addition, a gene (*ECM15 =* Anc_3.206) that is beside CEN3 in Saccharomycetales is instead found close to CEN2 in *H. uvarum* and *H. menglaensis* (Figure 3).

**Figure 3.**
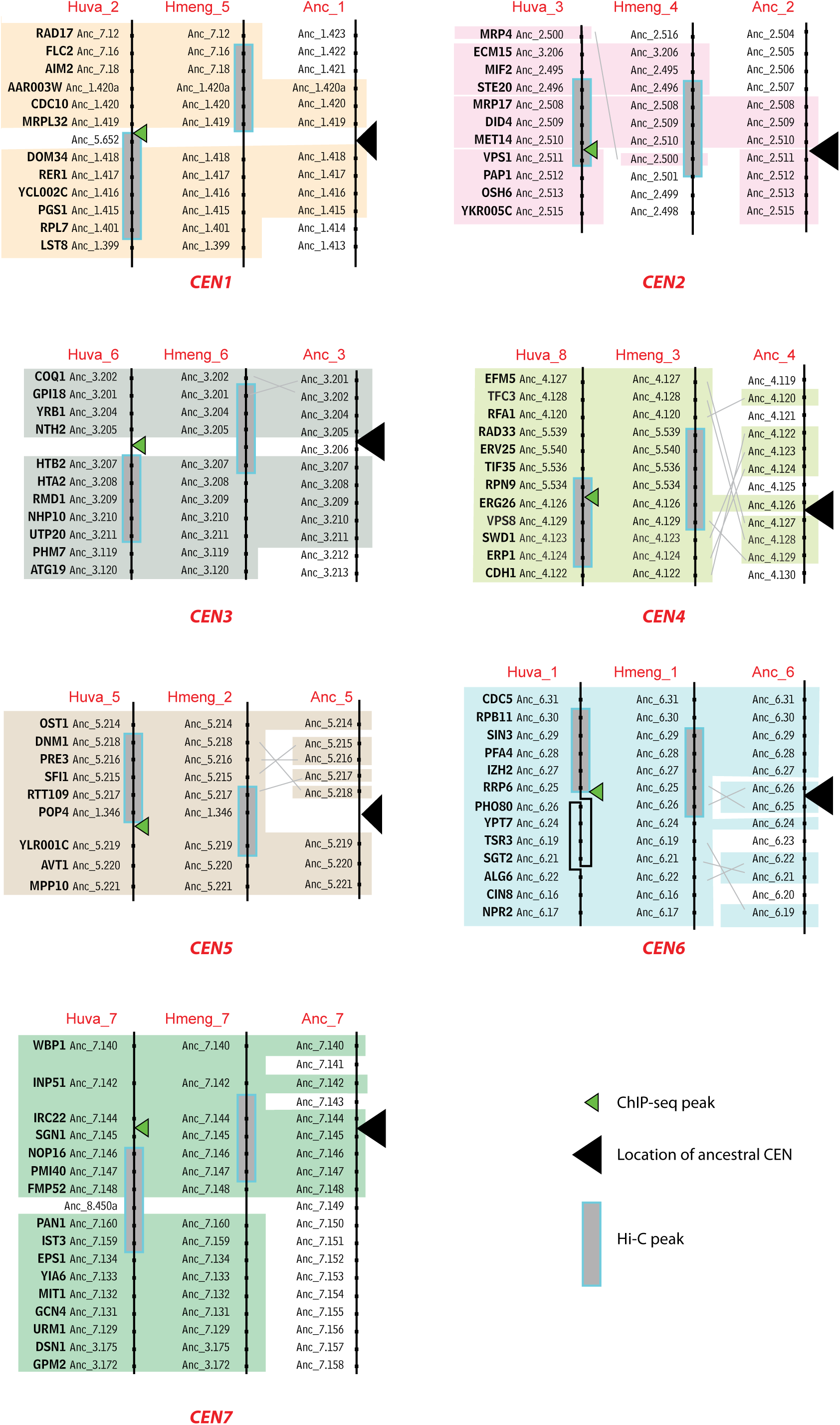
Conservation of gene order around centromeres in two *Hanseniaspora* species (Saccharomycodales) and the pre-WGD ancestor (Saccharomycetales). The panels compare gene order around each centromere in *H. uvarum* (Huva) and *H. menglaensis* (Hmeng) scaffolds, and the inferred gene order on the Ancestral (Anc) chromosomes of the Saccharomycetales pre-WGD ancestor [50]. Centromeres are named according to their ancestral chromosome number. Coloured backgrounds indicate regions of conserved gene order, and gray lines indicate rearrangements. At CEN6, there is an inversion in *H. uvarum* relative to *H. menglaensis*. *S. cerevisiae* gene names are shown for genes that are centromere-adjacent in at least two of the three genomes. “Anc” gene numbers are from Gordon et al. [50] and the Yeast Gene Order Browser (ygob.ucd.ie). ORFs that are not conserved between species are not shown.

### Centromere structure in Saccharomycodales

To explore the structure of *H. uvarum* centromeres, we used MEME 5.5.7 [52] to search for conserved motifs. We selected 500 bp surrounding the ChIP-seq peaks and looked for motifs that occurred zero or one times in each sequence. One clearly defined motif was present at every centromeric region (Figure 4A). The motif is 20 bp long and contains two copies of the sequence ACGCA that form a short inverted repeat, 5 nucleotides apart. Using the MEME suite program FIMO, we found that 8 of the 10 most significant matches to this motif in the *H. uvarum* genome are located at centromeres. The motif is very similar to one identified independently by Haase et al. [47], who called it the centromere-associated motif (CAM).

**Figure 4.**
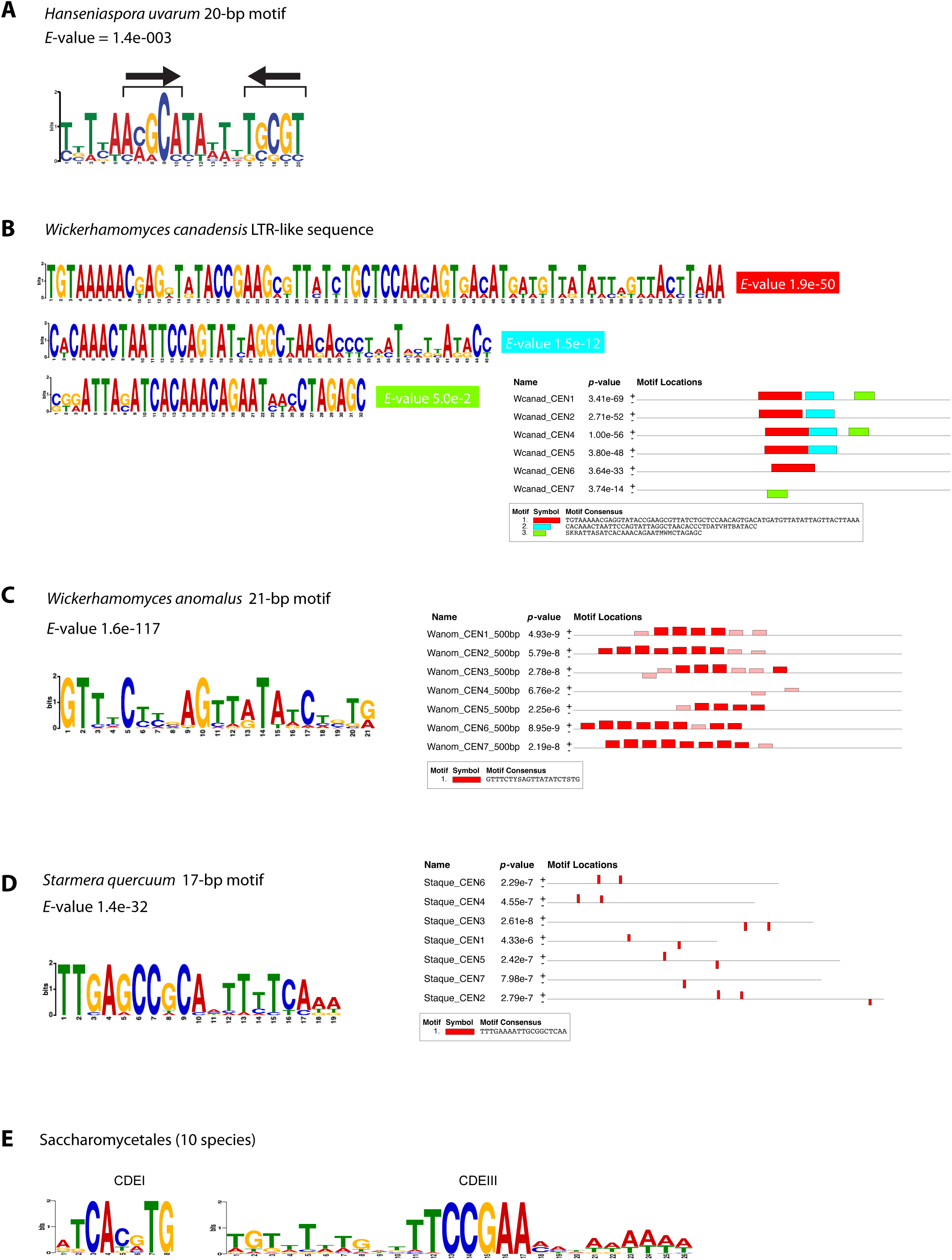
Centromere sequence motifs identified using MEME. **A.** *H. uvarum.* Sequence logo of the motif found at *H. uvarum* centromeres. The same motif was identified using MEME with either the OOPS (one occurrence per sequence) or ZOOPs (zero or one occurrence per sequence) options. The input sequences were 500 bp regions centered on the Cse4 ChIP-seq peaks. The arrows indicate a short inverted repeat. **B.** *W. canadensis*. The 3 most significant motifs identified in *W. canadensis* using MEME are shown with their locations within the input sequences, which were 500-bp regions (initially identified using BLASTN) within the Hi-C peaks. Together, the 3 motifs span a total of ∼200 bp at CEN1 and CEN4 and subsets of this sequence are present at 4 of the 5 other centromeres (it is absent at CEN3). MEME was run using the ANR (any number of repetitions) option, and a maximum motif length of 200 bp instead of the default value of 50 bp. **C.** *W. anomalus.* Left, the 21-bp motif identified by MEME (ANR option). Right, locations of matches to this motif in a 500-bp region from each centromeric region, chosen by centering on an initial candidate region identified by BLASTN. Red boxes show matches that were used by MEME to define the 21-bp motif, and pink boxes show additional (less significant) matches that were subsequently found by MEME by searching with the identified motif. **D.** *S. quercuum.* Left, the 17-bp motif identified by MEME (ANR option). Right, locations of matches to this motif in the candidate intergenic regions (1.2 – 2.4 kb long) at each Hi-C peak. **E.** Sequence logos of CDEI and CDEIII regions of point centromeres compiled from 10 Saccharomycetales species (including *S. cerevisiae*) for comparison (data from ref. [21]).

For *H. menglaensis*, most of the centromeric regions mapped by Hi-C include a large intergenic region (Figure S2), but our MEME analysis of these intergenic regions did not identify any significantly enriched motifs in them. Five of the *H. menglaensis* centromere regions are well conserved relative to *H. uvarum*, i.e. they are syntenic on both sides of the centromere, whereas there are some rearrangements at the other two (Figure 3). We extracted the sequences of the *H. menglaensis* intergenic regions corresponding to the locations of the *H. uvarum* ChIP-seq peaks for these five centromeres (CEN1, CEN3, CEN4, CEN5 and CEN7), which range in size from 1.1 kb to 4.2 kb, and searched for the presence of the *H. uvarum* motif using FIMO [52]. Matching motifs were found only at CEN4 and CEN7 (Figure S2). Thus, we have not identified any genomic feature that is shared by all the *H. menglaensis* centromeres and they appear to be featureless, whereas those of *H. uvarum* are sequence-defined. *Hanseniaspora* species are evolving rapidly [53], so it is perhaps not surprising that motifs are not well conserved between these two species.

### Centromere location in Phaffomycetales

Within Phaffomycetales, we found that gene order near centromeres is well conserved, particularly among *S. quercuum*, *W. anomalus*, *C. sargentensis* and *W. canadensis* (Figure 5). Gene order on both sides of CEN1, CEN2 and CEN4 is very similar in these four species, and there are only small changes at CEN5. For CEN3, CEN6 and CEN7, gene order is conserved on both sides of the centromere in three of these species, and the fourth shows conservation on one side. The three *Barnettozyma* species show a greater extent of rearrangement but conservation of centromere location is still clear, both within *Barnettozyma* and between it and the other Phaffomycetales. Genes that are centromere-proximal in one *Barnettozyma* species are usually also centromere-proximal in other *Barnettozyma* species, even where chromosome arm exchanges have occurred. All three *Barnettozyma* species share chromosome arm exchanges between one side of CEN4 and CEN1, and between the other side of CEN4 and CEN6, which occurred after *Barnettozyma* diverged from the other Phaffomycetales species (Figure 5).

**Figure 5.**
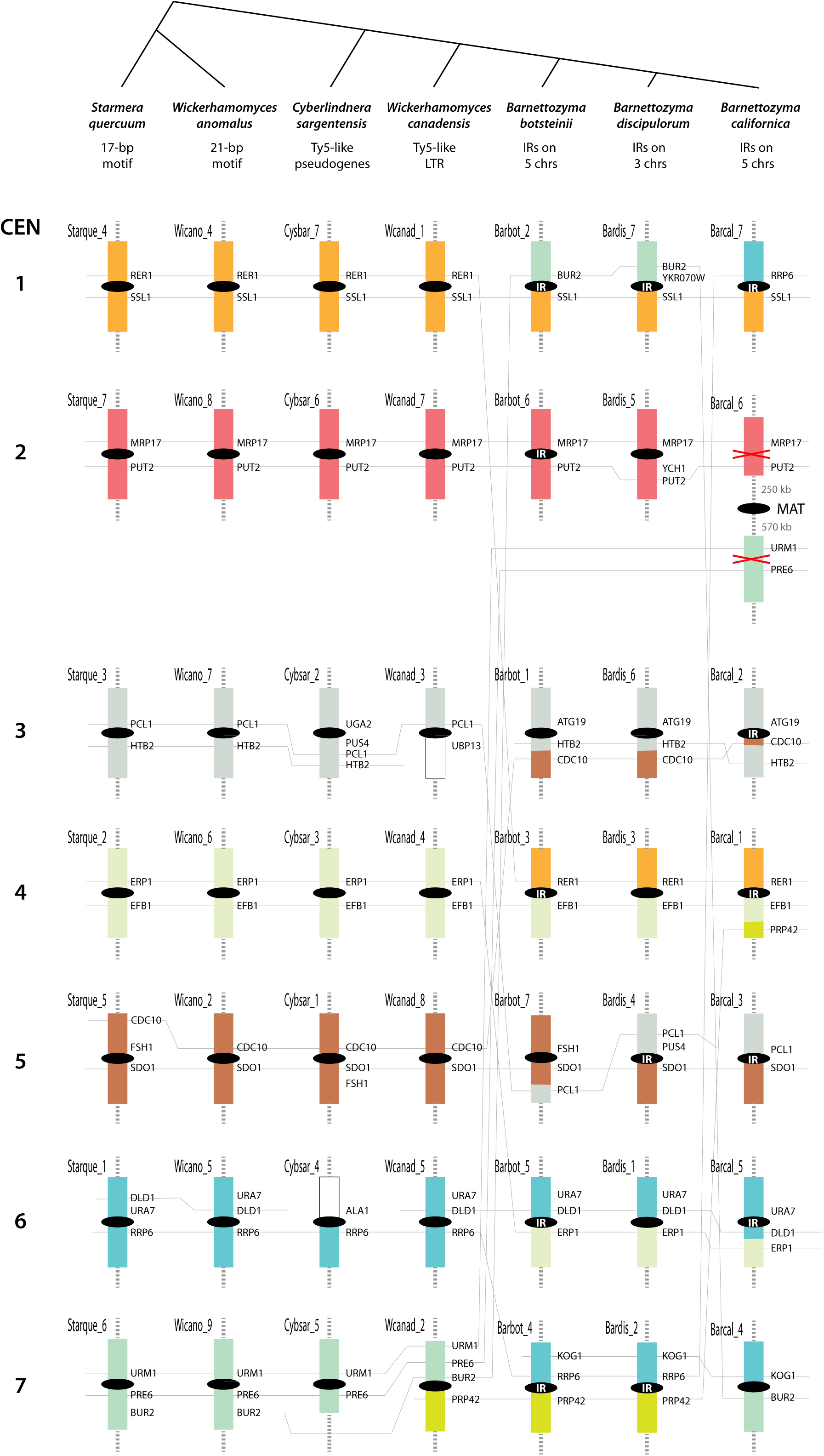
Conservation of centromere locations among Phaffomycetales species. Colored boxes represent regions of synteny conservation among the 7 species (gene content and order is mostly conserved, although there are some minor rearrangements within them). Only a small region near each centromere is shown; the remainder of each chromosome is indicated by whiskers. Some landmark *S. cerevisiae* gene names are shown to facilitate orientation with the detailed maps for each species in Supplementary Figures (Figs. S3, S4, S6, S8, S9, S10, S11). Gray lines connect orthologs in different species to highlight rearrangements. CEN numbers on the left indicate the ancestral centromere numbering scheme. Black ellipses represent centromeres, and “IR” indicates centromeres that contain inverted repeats.

There is less conservation of centromere locations between Phaffomycetales and either Saccharomycodales or Saccharomycetales, as expected because of the greater divergence time (approx. 208 versus 138 Myr; [13]), but conservation is still evident. Figure 6 compares the shared genes adjacent to centromeres of *Starmera quercuum* (representing Phaffomycetales), *H. uvarum* (representing Saccharomycodales), and the Anc_CENs of Saccharomycetales. For 6 of the 7 *S. quercuum* centromeres, 1-5 genes adjacent to it are also adjacent to centromeres in *H. uvarum* and the Saccharomycetales pre-WGD ancestor (Figure 6). The remaining centromere (CEN7) appears to have moved slightly in all Phaffomycetales but one centromere-adjacent gene (*URM1*) is shared between *S. quercuum* and *H. uvarum* (Figure 5; Figure 6).

**Figure 6.**
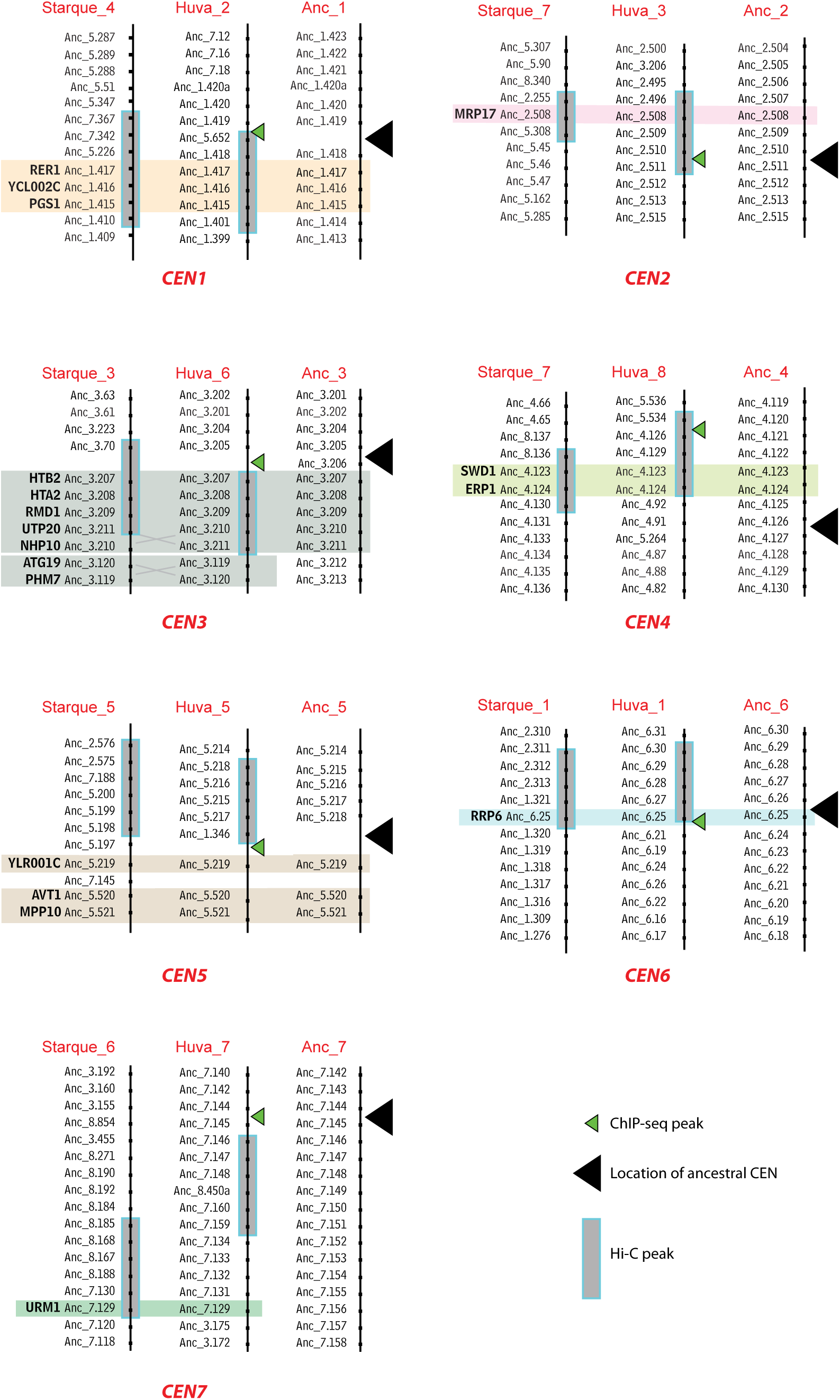
Genes conserved near the centromeres of Phaffomycetales, Saccharomycodales and Saccharomycetales. Colored boxes highlight genes that are present near centromeres in *S. quercuum* (Phaffomycetales), *H. uvarum* (Saccharomycodales) and the Saccharomycetales pre-WGD ancestor. The centromere number (e.g. CEN1) indicates the ancestral centromere name. Names beginning with Starque_ and Huva_ indicate chromosome or scaffold numbers in *S. quercuum* and *H. uvarum*, respectively. Some genes that lack homologs in other species have been omitted. *S. cerevisiae* gene names are shown for the conserved genes, and other genes are identified by using “Anc” gene numbers [50] which can be accessed via the Yeast Gene Order Browser (ygob.ucd.ie).

### Some centromeres have moved

Despite the overall conservation of centromere location, we found some examples of centromeres that have become relocated in individual Phaffomycetales species. Most strikingly, the location of CEN2 is almost completely conserved among six species but the equivalent location in *B. californica*, between the genes *MRP17* and *PUT2*, does not have a Hi-C signal (Figure 5). Instead, the centromere of this chromosome is located approximately 250 kb away. The new Hi-C centromere signal is at the Mating-Type (*MAT*) locus, as shown in Figure S3 and discussed further below. The same chromosome contains a region with genes *URM1* and *PRE6* that are adjacent to CEN7 in several of the other species. In *B. californica* CEN7 is on a different chromosome, between *BUR2* and *KOG1*, sharing synteny with one arm of CEN7 and one arm of CEN1 in *B. botsteinii* and *B. discipulorum* (Figure 5).

Some other centromeres have moved by smaller distances. For example, CEN3 lies between *PCL1* and *HTB2* in *S. quercuum* and *W. anomalus*, and beside *PCL1* in *W. canadensis*. However, in *C. sargentensis* CEN3 has moved by approximately 60 kb, and lies between *PUS4* and *UGA2* (Figure 5). At CEN1, the pericentric regions of both arms are shared by *S. quercuum*, *W. anomalus*, *C. sargentensis* and *W. canadensis*, but a rearrangement shared by all three *Barnettozyma* species has exchanged one arm with CEN4 (Figure 5). Closer examination of the gene order flanking CEN1 shows a series of rearrangement events (Figure 7). First, a block of four genes (8 kb) was inserted between *SSL1* and CEN1 in the common ancestor of *W. canadensis* and the three *Barnettozyma* species (yellow box in Figure 7). Next, the chromosome arm exchange with CEN4 occurred in the *Barnettozyma* common ancestor. Subsequently, in *B. discipulorum*, CEN1 moved by about 12 kb to its current location beside *YKR070W* (Figure 7). In summary, analysis of local gene order near centromeres in Phaffomycetales shows that rearrangements tend to be either small inversions or small insertion/deletions, or to be inter-chromosomal arm exchanges.

**Figure 7.**
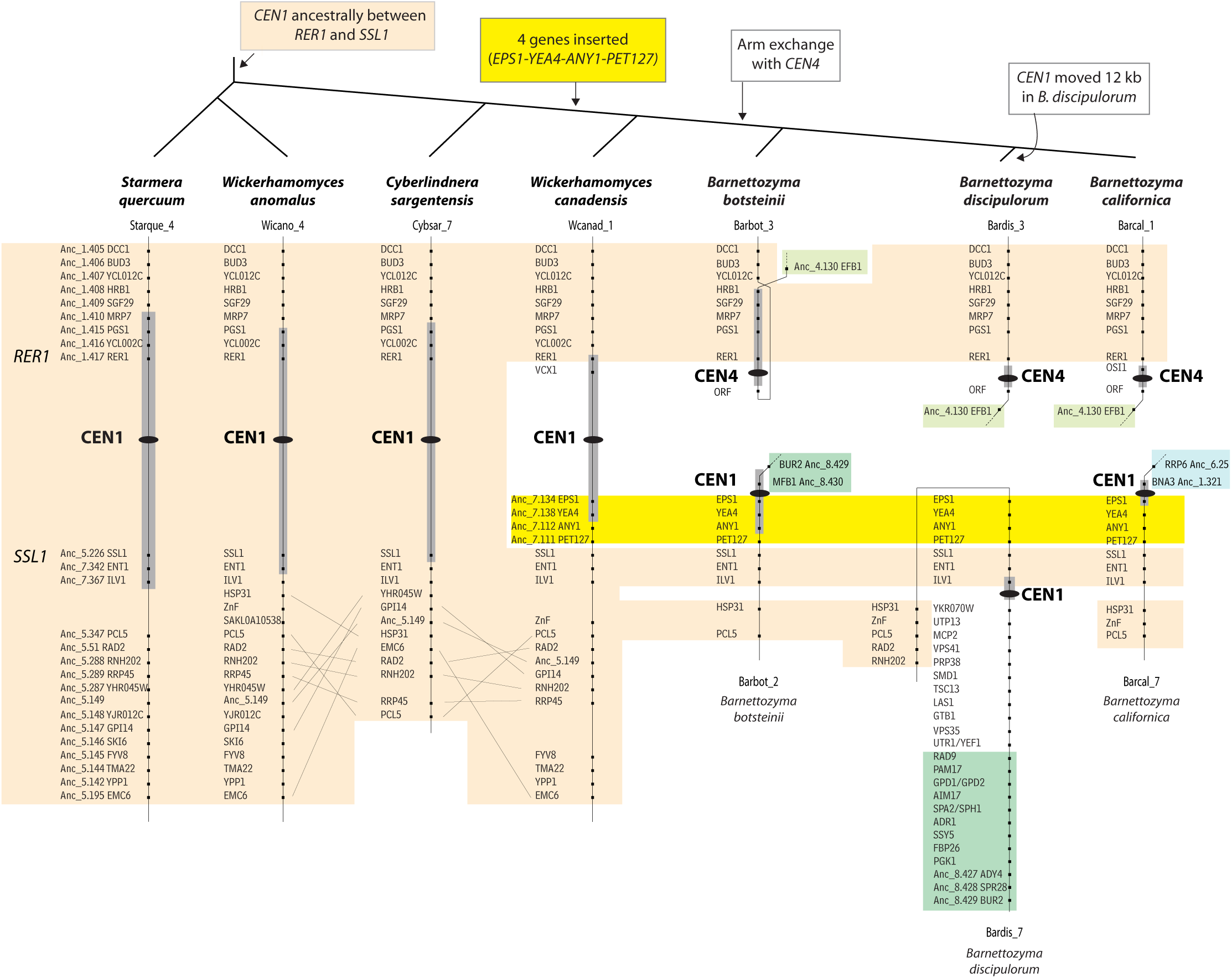
Details of rearrangements near CEN1 in Phaffomycetales. Black ellipses mark the inferred centromere locations, and gray rectangles around them indicate the extent of the 10-kb Hi-C peak region in each species (not drawn to scale). In the three outgroup species *S. quercuum*, *W. anomalus* and *C. sargentensis*, CEN1 is located between the genes *RER1* and *SSL1* and gene order in the centromere-proximal regions beyond these genes is relatively conserved (orange shading). In *W. canadensis*, the centromere is still between *RER1* and *SSL1* but a group of 4 genes (*EPS1, YEA4, ANY1* and *PET127*, highlighted in yellow) has been inserted between *SSL1* and the centromere. In the common ancestor of the three *Barnettozyma* species, an exchange of chromosome arms occurred so that *SSL1* and the 4 genes highlighted in yellow are near one centromere (designated CEN1), and *RER1* is near a different centromere (designated CEN4). CEN1 remains beside *EPS1* in *B. botsteinii* and *B. californica*, but a localized rearrangement in *B. discipulorum* has moved CEN1 by about 12 kb, placing it between *ILV1* and *YKR070W*.

### Centromere structure is highly variable in Cyberlindnera, Wickerhamomyces and Starmera (Phaffomycetales)

To characterize the structure of centromeres in Phaffomycetales, we looked for sequence motifs or shared structural characteristics at or near the Hi-C peaks on all chromosomes. The results for each species are described below and summarized at the top of Figure 5.

Two Phaffomyces species, *C. sargentensis* and *W. canadensis*, have centromere regions that contain sequences derived from retrotransposons. *C. sargentensis* has clusters of centromeric Ty5-like elements and LTRs at each of its Hi-C peak regions (Figure S4). All the copies of the element appear to be non-functional pseudogenes. There is a single cluster of Ty5-like sequences on every chromosome, in a large intergenic region, with >90% sequence identity between chromosomes in many cases. In dot-matrix comparisons of the *C. sargentensis* centromeric regions, we did not detect any sequences that are shared by all the centromere regions but not derived from Ty5-like elements (Figure S5). We therefore conclude that the centromeres of *C. sargentensis* must either consist of Ty5-like sequences, or are epigenetic with no conserved sequence (like *C. albicans*) but are a target that attracts integration of Ty5-like elements.

In *W. canadensis* (isolate CBS 1992) each Hi-C peak region contains a large intergenic region (2.2 - 9.3 kb; Figure S6). We identified a 200-bp sequence that is present, at least partially, in 6 of these 7 regions and is derived from a retrotransposon LTR (Figure 4B). This sequence occurs only in centromeric regions in the *W. canadensis* genome. We did not find any complete Ty5-like retrotransposons associated with this LTR (which we named LTR-Wc), but its identity as an LTR was confirmed by comparison to a related isolate (*Wickerhamomyces* sp. NRRL YB-2243) which has copies of LTR-Wc at other sites in the same centromere regions, including some that are longer (292 bp) and inserted cleanly into the LTRs of a different Ty5-like element (LTR-YB) that is absent from *W. canadensis* CBS 1992 (Figure S7). By MEME analysis, we did not find any motifs conserved among the *W. canadensis* CBS 1992 centromere regions, apart from LTR-Wc. *CEN3* does not contain LTR-Wc in either of the isolates, although it contains LTR-YB in *Wickerhamomyces* sp. NRRL YB-2243. Similar to *C. sargentensis*, we conclude that the centromeres of *W. canadensis* are either derived from the LTRs of Ty5-like element(s), or are epigenetic with no conserved sequence features.

In *W. anomalus*, there is a large intergenic region (3.7 - 7.9 kb) at each Hi-C peak (Figure S8). By BLASTN analysis we identified a sequence of ∼100 bp that is shared by 6 of these 7 intergenic regions (all except CEN4), and the CEN4 intergenic region contains a sequence that matches other parts of the CEN5 and CEN6 intergenic regions. Further analysis with MEME showed that the 6 *W. anomalus* loci contain an array of tandem copies of a 21-bp motif which we hypothesize is the centromere itself. Almost all the copies of the motif are on the same DNA strand, regularly spaced at 6 bp intervals (Figure 4C). There are 5-9 copies of this motif at each centromeric region except CEN4. The 21-bp motif is quite degenerate, with only 5 completely conserved positions (Figure 4C). By FIMO searches with its consensus sequence GTTTCTYSAGTTATATCTSTG we found that all 17 strong matches (FIMO *q* < 0.01) to this motif in the *W. anomalus* genome occur at centromeric regions, on 6 chromosomes. The CEN4 intergenic region contains two weaker matches to the motif (Figure 4C).

The isolate of *W. anomalus* (KG16) that we used for mapping the Hi-C data does not contain any Ty5-like elements or identifiable LTRs. However, we found that two other isolates of this species have Ty5-like elements in centromeric regions (Figure S8). Between them, they have one or more Ty5-like elements at every centromere, including some clusters, and the elements are inserted at different sites in each isolate, indicating recent mobility. The *W. anomalus* Ty5-like elements fall into three families: two (named Ty5.1 and Ty5.2) are autonomous elements coding for retrotransposon proteins with 28% amino acid sequence identity, and the third (named Ty5.L) is a non-autonomous large retrotransposon derivative (LARD) [31]. Ty5.L consists of two LTRs separated by 2.4 kb of noncoding DNA, which probably uses the proteins encoded by Ty5.1 or Ty5.2 for mobility. The LTRs of the three Ty5-like elements are substantially different from each other, but downstream of the LTR all three elements have a primer-binding site complementary to the anticodon stem of tRNA-iMet, which is diagnostic of the Ty5 family [54, 55]. The Ty5-like elements and their LTRs do not overlap with the 21-bp centromeric motif even though they are mostly located in the same intergenic regions (Figure S8). We therefore suggest that the *W. anomalus* centromere consists of the tandem iterations of the 21-bp motif, and that it attracts nearby integrations of at least three different types of Ty5-like element.

In *S. quercuum*, no repeats or sequence similarity among centromeric regions was detectable using BLASTN, and we did not find any Ty5-like elements or LTRs. However, MEME identified a 17-bp sequence motif that is present at least once in an intergenic region within the Hi-C peak on every chromosome (Figure 4D; Figure S9). Analysis of the whole *S. quercuum* genome sequence using FIMO showed that strong matches to this motif (FIMO *q* < 0.01) are present only in the Hi-C peak regions. The different copies of the motif are highly similar, and it includes the trimers CCG and CGC which are also present in the sequence-defined centromeres of *S. cerevisiae* and *H. uvarum* (Figure 4). We therefore infer that the *S. quercuum* centromeres are located at this 17-bp conserved motif.

The sequences of the 17-bp motif in *S. quercuum* and the 21-bp motif in *W. anomalus* do not appear to have any significant relationship to each other, nor to the LTRs present at centromeres in *C. sargentensis* and *W. canadensis*, which in turn are completely dissimilar even though they are both derived from retroelements in the Ty5 family. In contrast to the *Barnettozyma* species described in the next section, there are no large IRs at the centromeres of *Wickerhamomyces* or *Starmera*, and in *Cyberlindnera* the only IRs are those formed by parts of retrotransposons (Figure S5).

### Ongoing transition of centromere structure in the genus Barnettozyma (Phaffomycetales)

In the three *Barnettozyma* species that we investigated, all the Hi-C peaks coincide with long intergenic regions (Figure S3; Figure S10; Figure S11). Each species has IRs at the centromeres of 3-5 chromosomes, but not at their other centromeres (Figure 5). All the IRs consist of a pair of sequences in inverted orientation, with 99-100% sequence identity between the pair, ranging in size from 2.4-4.3 kb in *B. californica* to smaller IRs in *B. discipulorum* (1.0-1.4 kb) and *B. botsteinii* (0.5-0.8 kb). CEN1 is the only centromere that contains an IR in all three *Barnettozyma* species (Figure 5).

In *B. californica*, five centromeres contain IRs (Figure S3). The other two centromeres (CEN2 and CEN7) lack IRs but share a non-coding sequence, 460 bp long and 75% identical, that is not found anywhere else in the genome. CEN2 is the centromere that was newly formed at the *B. californica MAT* locus, after it diverged from the two other *Barnettozyma* species. It may therefore have been formed by duplication of the 460-bp sequence from CEN7. The 460-bp sequence at CEN2 is located in a 2.8-kb intergenic region between *MAT*a1 and *MAT*a2 genes on one side, and *MAT*α1 and *MAT*α2 genes on the other side (Figure S3).

Dot-matrix plots show that there is some sequence similarity among the IRs at different centromeres in *B. discipulorum*, but less in *B. botsteinii* and none in *B. californica* (Figure S12; Figure S13; Figure S14). This sequence similarity in *B. discipulorum* is primarily caused by the presence of tandem repeats of an octameric sequence motif (GTGGTTTT)_­_which is present at every *B. discipulorum* centromere, including those without IRs. By MEME analysis, we found that this motif is very strongly enriched at centromeres of *B. discipulorum* (*E-*value 3.0e-212) but not in the other two *Barnettozyma* species (Figure S15). FIMO searches using the sequence (GTGGTTTT)_­_as a query showed that 90 of the top 100 matches to this motif (FIMO *q* < 4.6e-5) in the *B. discipulorum* genome occur in the identified centromeric intergenic regions. This centromeric motif is 2 nucleotides different from the telomeric repeat of *B. discipulorum*, whose consensus is (GTGGGTGT)_­_, and many of the next-best matches are at telomeres. The motif is not centromere-specific in *B. californica* or *B. botsteinii*.

We conclude that centromere structure in the genus *Barnettozyma* is polymorphic – some centromeres have IRs and others do not – and that a transition of structure is underway in this genus. The fact that the new centromere at the *B. californica MAT* locus does not contain an IR, and was probably formed by duplicating the IR-lacking centromere CEN7, suggests that the direction of the structural transition in *Barnettozyma* is away from IR-containing centromeres and toward centromeres that are either featureless, or (in *B. discipulorum*) based on a short sequence motif.

## Discussion

Centromere structures were known to have diversified greatly during the ∼404 Myr evolutionary span of the subphylum Saccharomycotina (reviewed in [5]). Our results now show that diversification has continued to occur on shorter and more recent timescales, such as the ∼144 Myr divergence time among the Phaffomycetales species we compared [13]. Remarkably, most of this evolutionary reorganization of centromere structure has occurred in the absence of major changes of centromere location. For example, ancestral CEN5 has many different structures in different species, yet it has retained significant local synteny (Figure 5; Figure 6). Despite their variation in structure, we found that almost all centromeres are conserved in location among Phaffomycetales species, with only minor rearrangements, and that there is extensive synteny conservation among the 7 centromeres of Phaffomycetales, the 7 centromeres of Saccharomycodales, and 7 of the 8 ancestral centromeres of Saccharomycetales.

It is difficult to tell whether a new centromere structure arose *in situ* by mutation of the previous centromere itself, or by the formation of a new centromere at a site close to the previous location but separate from it. Helsen et al. [51] recently reported that transitions between different types of point centromeres with characteristically different CDEII region lengths are ongoing in some Saccharomycetales species, with CDEII length being polymorphic in some species including *S. cerevisiae*; these are examples of a small change of a centromere’s structure *in situ* by mutation. In *Naumovozyma* species (Saccharomycetales) new point centromeres with substantially different CDE motif sequences arose either at, or very close to, the ancestral centromere locations of Saccharomycetales [22]. The situation that we and Haase et al. [47] found in Saccharomycodales species is similar to that in *Naumovozyma*: the sequences of the sequence-defined centromeres of Saccharomycodales are different from that of Saccharomycetales, so either they changed *in situ* by mutation, or new centromeres emerged beside old centromeres, but it is difficult to distinguish between these two possibilities. In *Barnettozyma* we found an example (CEN4 in *B. californica* and *B. discipulorum*) where a centromere’s location has been conserved between two species, but one species has an IR and the other does not, without any changes in the local gene order except for the presence of an extra gene that forms part of the IR. This example suggests that IR-containing and IR-lacking centromeres can be interchanged *in situ* by structural mutations that create or destroy an IR, without a change of location.

Evolutionary emergence of new centromeres near the old centromeres is likely to be much better tolerated than emergence of new centromeres at distant locations, because any change of the centromere structure in a species must proceed via an intermediate polymorphic stage during which different alleles of a centromere need to be able to function together when heterozygous in the same nucleus. The genus *Barnettozyma* appears to be going through this polymorphic stage, because IR-containing and IR-lacking centromeres coexist in each species. Yeast centromeres remain close together in space during most of cell division and are located at the nuclear periphery [44]. It has been proposed that binding of CENP-A (Cse4) to these centromere clusters results in the formation of a cloud of free CENP-A molecules, which can then occupy DNA regions in the same space [56]. This cloud may explain why new epigenetic centromeres (neocentromeres) form close to previous centromere locations in *C. albicans* and *C. parapsilosis* [5, 32]. The CENP-A cloud may select for conservation of the location of a centromere, while allowing divergence of its structure.

Of the 49 Phaffomycetales centromeres that we mapped (7 species x 7 chromosomes each), only one – the *MAT* locus centromere of *B. californica* – showed long-distance relocation to a previously non-centromeric site. On this chromosome, a centromere was lost from the *MRP17-PUT2* locus and a centromere was gained at the *MAT* locus, 250 kb away (Figure 5). This relocation event occurred in the absence of any major chromosomal rearrangements, because in all three *Barnettozyma* species the *MAT* locus and *MRP17-PUT2* are on the same chromosome and about 250 kb apart; the former is the centromere in *B. californica*, and the latter is the centromere in *B. discipulorum* and *B. botsteinii*. All three *Barnettozyma* species appear to be homothallic, in that both *MAT***a** and *MAT*α genes are present at their *MAT* loci. It is possible that in *B. californica*, proximity to the centromere may control expression of one or other set of *MAT* genes, similar to centromere-dependent repression of the *MAT* locus in *Ogataea polymorpha* [37], enabling each cell to be functionally *MAT***a** or *MAT*α. There may therefore have been a selective advantage in moving the centromere to the *MAT* locus in *B. californica*, consistent with the evolutionary trend toward homothallism in budding yeasts [57].

Our results suggest that sequence-defined centromeres have emerged independently in several separate yeast lineages. Τhe point-like centromeres of Saccharomycetales and Saccharomycodales are older than previously thought, and it has recently been hypothesized that they share a common ancestor related to the LTR of a Ty5-like retroelement [47]. The centromere structures we found in *W. canadensis* and *C. sargentensis* are consistent with this hypothesis. However, the shorter centromere sequence motifs that we identified in *S. quercuum* (17 bp) and *W. anomalus* (21 bp) have no obvious similarity to Ty5 LTRs, nor to each other, so they appear to have a different origin. Importantly, in *W. anomalus* we found that Ty5-like elements are present in two isolates of this species but absent in a third, which is more consistent with the hypothesis that Ty5-like elements are centromere-associated, than that they are centromere-making. It is therefore unclear to us whether an origin of sequence-defined centromeres from the LTRs of a Ty5-like element, as was hypothesized [47] for the common ancestor of Saccharomycodales and Saccharomycetales, can be extrapolated further back to their common ancestor with Phaffomycetales as well. Furthermore, our results from *Barnettozyma* species indicate that some of their centromeres are defined by IRs, with no connection to Ty5-like elements or LTRs, unless this relationship has been completely obscured by sequence divergence.

The diversity of structures makes it difficult to deduce the ancestral form of centromeres in budding yeasts, or the direction of change when one structure replaces another, but it is evident that transitions in structure have occurred frequently during the evolution of yeasts and even within the single genus *Barnettozyma*. We speculate that structural transitions may be the result of yeast cells sporulating in nutrient-poor conditions in which they only have sufficient resources to make one spore instead of four, leading to centromere drive – competition between the alleles of a centromere for inclusion in the spore [58–61].

## Methods

### Yeast strains and genome data

Details of yeast isolates used for genome sequence analysis and for Hi-C and ChIP-seq data generation, with NCBI accession numbers, are given in Table S1. We generated new highly contiguous genome sequence assemblies for four species, using a combined Oxford Nanopore Technologies (ONT) long-read and Illumina short-read approach. Genomes were annotated using YGAP [62].

For *S. quercuum* CBS 2283, genomic DNA was extracted using a MasterPure Yeast DNA purification kit (BioLife Technologies). Libraries were generated using Native Barcoding Kit 24 from ONT (SQK-NBD112.24) and sequenced on a FLOMIN-106 flow cell (R9.4.1) using a MinION MK1C. Basecalling and demultiplexing was carried out on the MK1C using MinKNOW (v21.11.6) with default settings. Sequencing reads were filtered using NanoFilt v2.8 to retain those >Q7 and >1 kb (76,013 reads), assembled using Canu 2.2 [63], and polished with short read data (from NCBI accession SRR6475995) using 5 rounds of NextPolish 1.4 [64].

For *W. canadensis* CBS 1992, genomic DNA was extracted using a QIAGEN genomic tip 100/G. Libraries were generated using a Ligation sequencing kit (SQK-LSK109) and sequenced on a FLOMIN-106 flow cell (R9.4.1) using a MinION MK1C. Base-calling was performed using Guppy (v.4.2.2+effbaf8). Sequencing reads were filtered using NanoFilt to retain those >Q10 and >10 kb (313,617 reads), assembled using Canu v2.2 and polished with short read data (SRR6475885) using 5 rounds of NextPolish 1.4. Two contigs were subsequently joined together using data from an assembly of the short reads only.

For *B. californica* UCD09, genomic DNA was extracted using a QIAGEN genomic tip 100/G. Libraries were generated using a Rapid Barcoding Sequencing Kit (SQK-RBK004) and sequenced on a FLOMIN-106 flow cell (R9.4.1) using a MinION MK1C. Base-calling was performed using Guppy (v.4.0.9) in MInKNOWv4.0.13. Sequencing reads were filtered using NanoFilt v2.3 to retain those >Q8 and >1 kb and were assembled using Canu v1.8 using one round of polishing with short read data [65] with Pilon v1.23 [66].

For *B. botsteinii* CBS 16679, genomic DNA was isolated using a MasterPure Yeast DNA purification kit. Libraries were generated using Native Barcoding Kit 24 from ONT (SQK-NBD112.24) and sequenced on a FLOMIN-106 flow cell (R9.4.1) using a MinION MK1C. Basecalling and demultiplexing was carried out on the MK1C using MinKNOW (v21.11.6) with default settings. Reads were filtered using NanoFilt to retain those >Q7 and >5 kb (86,523 reads), assembled using Canu V 2.2 and polished with short read data from the Hi-C analysis using 5 rounds with NextPolish 1.4.

We recently isolated *B. discipulorum* UCD2008 as a new species in the genus *Barnettozyma*. A publication reporting its genome sequence is currently under review [38].

### Hi-C data generation and mapping

For crosslinking, yeast strains were grown overnight in 5 mL of YPD media. Cells were spun down and resuspended in 10 mL of 1% formaldehyde and incubated at room temperature for 20 mins with periodic vortexing. Cells were quenched with glycine (125 mM final concentration) for 15 min at room temperature with periodic vortexing to end the cross-linking. Cells were spun down (1000 x *g*) for 1 min and washed using PBS. Cells were then resuspended in 400 µL of PBS and shipped to Phase Genomics (Seattle, WA, USA) for Hi-C sequencing. Hi-C reads were aligned to the relevant genome assemblies with BWA MEM v0.7.17-r1188 [67]. Each member of the paired end reads was aligned separately. Reads with a maximum BWA MAPQ score of 60 (i.e. the BWA aligner’s highest probability that the mapped position reported is correct) were retained, ignoring mismatches and truncation. An interaction was recorded when each read in a pair mapped with a BWA MAPQ score of 60. Hi-C interaction heatmaps were created for all genomes, with one pixel equivalent to a 10 kb window. To more accurately identify the most likely centromere regions, the number of interactions between one chromosome and all other chromosomes in a species was calculated in 10 kb windows, stepping the windows by 1 kb (*trans* interactions). The mean and standard deviation of the interactions of each 10 kb window was calculated and used to derive the IoD (StandardDeviation^2^ / Mean). For *cis* interactions, the IoD was calculated after first removing the within-chromosome interactions on the diagonal.

### Expressing H. uvarum CSE4 gene with 3 x HA tag

A DNA fragment was synthesized containing the *H. uvarum CSE4* gene with its native promoter, with a 3x HA (haemagglutinin) tag inserted between amino acids 34 and 35 (Figure S1), and was cloned into plasmid pJJH3200 [46]. The resulting plasmid pJJH3200_CS4E_HA was transformed into *H.uvarum* strain HHO1 (a derivative of DSM 2768) by electroporation as described [46]. Expression of the CSE4_HA tagged protein was confirmed by western blot analysis, using the mouse epitope tag antibody Anti-HA.11 (BioLegend 901516), at a 1:2000 dilution in 20 mL of 5% skim milk powder in PBS+tween 0.05% and HRP-conjugated secondary antibody anti-mouse IgG (Cell Signaling Technology 7076P2) at 1:10000 dilution. Immunoblots were visualized using the Pierce ECL western blotting substrate (Thermo Fisher Scientific 32106) and enhanced chemiluminescence (Vilber Fusion FX spectra).

### ChIP-seq analysis of Cse4 binding sites in H. uvarum

Chromatin immunoprecipitation was performed as described [34]. HHO1 strain with pJJH3200_CS4E_HA plasmid was grown in 200 mL YPD + 300 μg/mL hygromycin B until reaching log phase, after which crosslinking was initiated using 1% formaldehyde at room temperature. The reaction was quenched by adding 2.5 M glycine. Crosslinked cells were then washed with TBS buffer (100 mM Tris–HCl, pH 7.5, 150 mM NaCl) and resuspended in FA lysis buffer (50 mM HEPES, pH 7.5, 150 mM NaCl, 1 mM EDTA, 1% Triton X-100, 0.1% sodium deoxycholate, 0.1% SDS) supplemented with 1 mM PMSF. Cell lysis was carried out using glass beads, and the chromatin was sheared by sonication. For immunoprecipitation, fragmented chromatin from three biological replicates was incubated with EZview Red Anti-HA Affinity Gel (Sigma–Aldrich). A separate control sample was treated with Mouse IgG1 isotype control (Cell Signaling Technology). Following several wash steps, the DNA-protein complexes were eluted using HA peptide (Sigma–Aldrich). Crosslinks were reversed, and DNA was purified by phenol–chloroform extraction prior to sequencing. Sequencing was performed by Novogene. ChIP-seq reads were aligned to the genome assembly of *H. uvarum* isolate CBA6001 with BWA MEM v0.7.17-r1188 [67] after filtering by Skewer v0.2.2 [68].

## Acknowledgements

We thank Adam Ryan, Eoin Ó Cinnéide, Letal Salzberg, Sean Bergin, Hawraa A. A. Almotawaa, Ben O’Leary Chaney, and Cristina Gonzales for assistance with genome sequencing and assembly. We thank Prof. Dr. Jürgen Heinisch (Universität Osnabrück, Germany) for the plasmid and strain used for *H. uvarum* ChIP-seq. This work was supported by Research Ireland (grant numbers 19/FFP/6668, 20/FFP-A/8795 and IRCLA/2023/1951) and the European Research Council (789341).

## Author contributions

C.H. carried out the ChIP-seq experiments, and DNA isolation for Hi-C sequencing of all species. K.P.B. mapped and analyzed the Hi-C and ChIP-seq data. G.B. and K.H.W. supervised the project and assisted with data analysis and writing.

## Supplementary Figures

**Figure S1. Construction of an *H. uvarum* strain expressing an HA-tagged Cse4 protein for ChIP-seq analysis.**

**A**. A synthetic gene coding for HuCse4 with 3 HA tags between codons 34 and 35 was cloned into plasmid pJJH3200 [46] and transformed into *H. uvarum* strain HHO1 by electroporation.

**B**. Expression of HuCse4-HA in *H. uvarum* was detected by Western blot using anti-HA antibody. Lane L: Precision Plus Protein Dual Color Standards (Bio-RAD). Sizes are shown in kD. Lane C: Protein extract from untransformed *H. uvarum* HHO1. Lane S: Protein extract from transformed *H. uvarum* HHO1 strain expressing pJJH3200_CSE4_HA plasmid.

**C.** Multiple sequence alignment of Cse4 proteins from five *Hanseniaspora* species, showing the site at which the 3xHA tag was inserted in *H. uvarum* Cse4.

**Figure S2. Locations of centromeres in *H. menglaensis* by Hi-C mapping.** Yellow boxes show the locations of the 10-kb Hi-C peak windows identified on each scaffold, with the start position (kb) of each window labeled on its left. Blue arrows show annotated genes, named according to their *S. cerevisiae* orthologs and ancestral (Anc) gene numbers where possible. The location of motifs at CEN4 and CEN7 matching those found at *H. uvarum* centromeres are shown. The CEN numbers reflect the ancestral centromere numbering that has been applied to all species. The Scaffold numbers are specific to the *H. menglaensis* (Hmeng) assembly.

**Figure S3. Location of centromeres in *B. californica* by Hi-C mapping**. Yellow boxes show the locations of the 10-kb Hi-C peak windows identified on each scaffold, with the start position (kb) of each window labeled on its left. Magenta arrows show the location of large Inverted Repeats (IRs), with their length and percent sequence identity shown. The red boxes indicate a ∼460 bp sequence shared by CEN7 (Scaffold 4) and CEN2 (Scaffold 6). Blue arrows show annotated genes, named according to their *S. cerevisiae* orthologs and ancestral (Anc) gene numbers where possible. The CEN numbers reflect the ancestral centromere numbering that has been applied to all species. The Scaffold numbers are specific to the *B. californica* (Barcal) assembly. Genes whose names are underlined were used as landmarks in Figure 5; other landmark genes that lie outside the illustrated region are named at the edges, with the number of ORFs separating them shown in parentheses.

**Figure S4. Location of centromeres in *C. sargentensis* by Hi-C mapping.** Yellow boxes show the locations of the 10-kb Hi-C peak windows identified on each scaffold, with the start position (kb) of each window labeled on its left. Magenta arrows show sequences annotated as Ty5-like pseudogenes. Blue arrows show annotated genes, named according to their *S. cerevisiae* orthologs and ancestral (Anc) gene numbers where possible. The CEN numbers reflect the ancestral centromere numbering that has been applied to all species. The Scaffold numbers are specific to the *C. sargentensis* (Cybsar) assembly. Genes whose names are underlined were used as landmarks in Figure 5; other landmark genes that lie outside the illustrated region are named at the edges, with the number of ORFs separating them shown in parentheses.

**Figure S5. Dot matrix plot comparing centromere regions in *C. sargentensis.*** The sequences of the longest intergenic region within the Hi-C peaks on each chromosome were extracted and concatenated. The plot was generated using DNAMAN with a threshold of 17 mismatches per 56-bp window.

**Figure S6. Location of centromeres in *W. canadensis* by Hi-C mapping.** Yellow boxes show the locations of the 10-kb Hi-C peak windows identified on each scaffold, with the start position (kb) of each window labeled on its left. Magenta boxes show sequences derived from LTR-Wc (see Figure 4B for detail), which are present at every centromere except CEN3. Blue arrows show annotated genes, named according to their *S. cerevisiae* orthologs and ancestral (Anc) gene numbers where possible. The CEN numbers reflect the ancestral centromere numbering that has been applied to all species. The Scaffold numbers are specific to the *W. canadensis* (Wcanad) assembly. Genes whose names are underlined were used as landmarks in Figure 5; other landmark genes that lie outside the illustrated region are named at the edges, with the number of ORFs separating them shown in parentheses.

**Figure S7. LTRs in *Wickerhamomyces* species NRRL YB-2243.** Examples of LTR-containing regions from CEN5, CEN6 and CEN3 are shown. At CEN5 and CEN6, copies of LTR-Wc have become inserted into LTR-YB. LTR-Wc has sequence similarity to the 200-bp motif identified in centromeres of *W. canadensis*. LTR-YB is present in *Wickerhamomyces* sp. NRRL YB-2243 but not in *W. canadensis* CBS 1992.

**Figure S8. Location of centromeres in *W. anomalus* by Hi-C mapping**. Yellow boxes show the locations of the 10-kb Hi-C peak windows identified on each scaffold, with the start position (kb) of each window labeled on its left. Magenta boxes show sequence regions that were initially identified by BLASTN as being similar among centromeres, and which contain multiple iterations of a 21-bp motif (Figure 4C). The gene maps and coordinates are from the assembly of the primary haplotype of *W. anomalus* isolate KG16, which does not contain any identified Ty5-like elements or LTRs. Two other isolates of *W. anomalus*, KCTC 27761 (green) and CpEC_Uees (magenta) contain three types of Ty5-like element (labeled Ty5.1, Ty5.2 and Ty5.L) integrated at different sites as shown. NCBI accession numbers are shown for each isolate. Blue arrows show annotated genes, named according to their *S. cerevisiae* orthologs and ancestral (Anc) gene numbers where possible. The CEN numbers reflect the ancestral centromere numbering that has been applied to all species. The Scaffold numbers are specific to the *W. anomalus* (Wanom) KG16 assembly. Genes whose names are underlined were used as landmarks in Figure 5; other landmark genes that lie outside the illustrated region are named at the edges, with the number of ORFs separating them shown in parentheses.

**Figure S9. Location of centromeres in *S. quercuum* by Hi-C mapping.** Yellow boxes show the locations of the 10-kb Hi-C peak windows identified on each scaffold, with the start position (kb) of each window labeled on its left. Magenta boxes show occurrences of the 17-bp motif identified in Figure 4D. Blue arrows show annotated genes, named according to their *S. cerevisiae* orthologs and ancestral (Anc) gene numbers where possible. The CEN numbers reflect the ancestral centromere numbering that has been applied to all species. The Scaffold numbers are specific to the *S. quercuum* (Staque) assembly. Genes whose names are underlined were used as landmarks in Figure 5; other landmark genes that lie outside the illustrated region are named at the edges, with the number of ORFs separating them shown in parentheses.

**Figure S10. Location of centromeres in *B. discipulorum* by Hi-C mapping**. Yellow boxes show the locations of the 10-kb Hi-C peak windows identified on each scaffold, with the start position (kb) of each window labeled on its left. Magenta arrows show the location of large Inverted Repeats (IRs), with their length and percent sequence identity shown. Blue arrows show annotated genes, named according to their *S. cerevisiae* orthologs and ancestral (Anc) gene numbers where possible. The CEN numbers reflect the ancestral centromere numbering that has been applied to all species. The Scaffold numbers are specific to the *B. discipulorum* (Bardis) assembly. Genes whose names are underlined were used as landmarks in Figure 5; other landmark genes that lie outside the illustrated region are named at the edges, with the number of ORFs separating them shown in parentheses.

**Figure S11. Location of centromeres in *B. botsteinii* by Hi-C mapping**. Yellow boxes show the locations of the 10-kb Hi-C peak windows identified on each scaffold, with the start position (kb) of each window labeled on its left. Magenta arrows show the location of large Inverted Repeats (IRs), with their length and percent sequence identity shown. Blue arrows show annotated genes, named according to their *S. cerevisiae* orthologs and ancestral (Anc) gene numbers where possible. The CEN numbers reflect the ancestral centromere numbering that has been applied to all species. The Scaffold numbers are specific to the *B. botsteinii* (Barbot) assembly. Genes whose names are underlined were used as landmarks in Figure 5; other landmark genes that lie outside the illustrated region are named at the edges, with the number of ORFs separating them shown in parentheses.

**Figure S12. Dot matrix plot comparing centromere regions in *B. discipulorum.*** The sequences of the longest intergenic region within the Hi-C peaks on each chromosome were extracted and concatenated. The plot was generated using DNAMAN with a threshold of 17 mismatches per 56-bp window.

**Figure S13. Dot matrix plot comparing centromere regions in *B. botsteinii.*** The sequences of the longest intergenic region within the Hi-C peaks on each chromosome were extracted and concatenated. The plot was generated using DNAMAN with a threshold of 17 mismatches per 56-bp window.

**Figure S14. Dot matrix plot comparing centromere regions in *B. californica.*** The sequences of the longest intergenic region within the Hi-C peaks on each chromosome were extracted and concatenated. The plot was generated using DNAMAN with a threshold of 17 mismatches per 56-bp window.

**Figure S15. MEME analysis of motifs enriched at centromeres in *Barnettozyma* species.** For each species, the input sequences were the largest intergenic region at each Hi-C peak. MEME was run using default parameters and the ANR (“any number of repeats”) option, and identified the three most significantly enriched motifs. The locations of the motifs, relative to the Inverted Repeats (black arrows) present at some centromeres, are shown. In *B. discipulorum*, Motifs 1 consists of tandem iterations of an 8-mer with the consensus sequence GTGGTTTT, and occurs on all chromosomes. Motif 2 is a variant of Motif 1.

## List of Supplementary Tables

**Table S1.** Sources of genome sequence data, Hi-C data, and ChIP-seq data used in this study.

**Table S2.** Genome assembly statistics for yeast newly sequenced in this study.

**Table S3**. Location of centromeres mapped by Hi-C in each species. For each scaffold, the number shown is the start position (in kb) of the 10-kb Hi-C peak window identified.

